# Analysing the effect of full-length and C-terminally truncated progranulin on proliferation, colony formation, and migration in HepG2 and U87 cells

**DOI:** 10.1101/2025.08.01.668100

**Authors:** Alexander M. Hofer, Luisa Tobler, Marc-David Ruepp, Oliver Mühlemann

## Abstract

Progranulin, the precursor protein to seven and a half distinct granulin motifs (GRNs), has been implicated in a broad range of diseases. Progranulin depletion is one of the most frequent causes for hereditary Frontotemporal Dementia (FTD). On the other hand, elevated progranulin levels have been associated with increased malignancy of many tumours, manifesting in increased cell proliferation, migration, metastasis formation, and reduced sensitivity to chemotherapeutics. While some functions can be unambiguously attributed to either full-length progranulin or one or multiple of the different GRNs, much about the interplay between progranulin and GRNs remains unknown. Here, we aimed to test the effect of progranulin overexpression on cell-based tumorigenicity assays, assessing proliferation, migration, and colony formation, using the hepatocellular carcinoma cell line HepG2 and the glioblastoma cell line U87. We transduced these cells with lentiviral vectors to overexpress full-length progranulin, two different C-terminally truncated progranulin proteins, lacking either the last two or the last four GRNs, or a triple FLAG-tagged maltose binding protein as a control. We observed increased colony formation in HepG2 overexpressing the full-length progranulin but not the C-terminally truncated constructs. The U87 cell lines were neither affected by an increase in progranulin levels nor by the depletion of progranulin.

## Introduction

The secreted glycoprotein progranulin (PGRN) is expressed ubiquitously throughout human tissues and has been associated with different functions, ranging from early development ^1^, regulating inflammation ^2,3^, cancer ^4,5^ to neuroprotection ^6–8^. PGRN is encoded by the granulin gene (*GRN*), located on chromosome 17q21.32 ^9^, and consists of seven and a half cysteine-rich motifs called granulins (GRNs). PGRN is cleaved by various proteases, which proteolytically cut in the linker regions, thereby releasing individual GRNs and oligo-GRNs ^2,10–12^.

The functional relationship between individual GRNs, oligo-GRNs, and full-length PGRN remains largely elusive, with the full-length PGRN being more studied than individual GRNs. PGRN has been shown to act as an anti-inflammatory agent, competing with tumour necrosis factor alpha (TNFα) for the binding of tumour necrosis factor receptor (TNFR) 1 & 2 ^3^. In addition, the inhibition or knockout of the secretory leukocyte protease inhibitor (SLPI), an anti-inflammatory protein, resulted in increased processing of PGRN into GRNs, ultimately resulting in elevated inflammation, which can be rescued through the addition of full-length PGRN but not by the addition of GRNs ^2,3^. Tang and colleagues determined the minimal fraction of PGRN needed for binding to TNFRs and for competing with TNFα. This resulted in the engineering of Atsttrin (Antagonist of TNFα/TNFR signalling via Targeting to TNF Receptors), which consists of portions of GRNs F, A, and C, as well as the linker regions between them (order of granulin motifs depicted in Figure 1A). Atsttrin has since been studied as a potential therapeutic for inflammatory arthritis ^3^. It is generally thought that PGRN is anti-inflammatory, while certain GRNs can act as pro-inflammatory cues.

**Figure 1:**
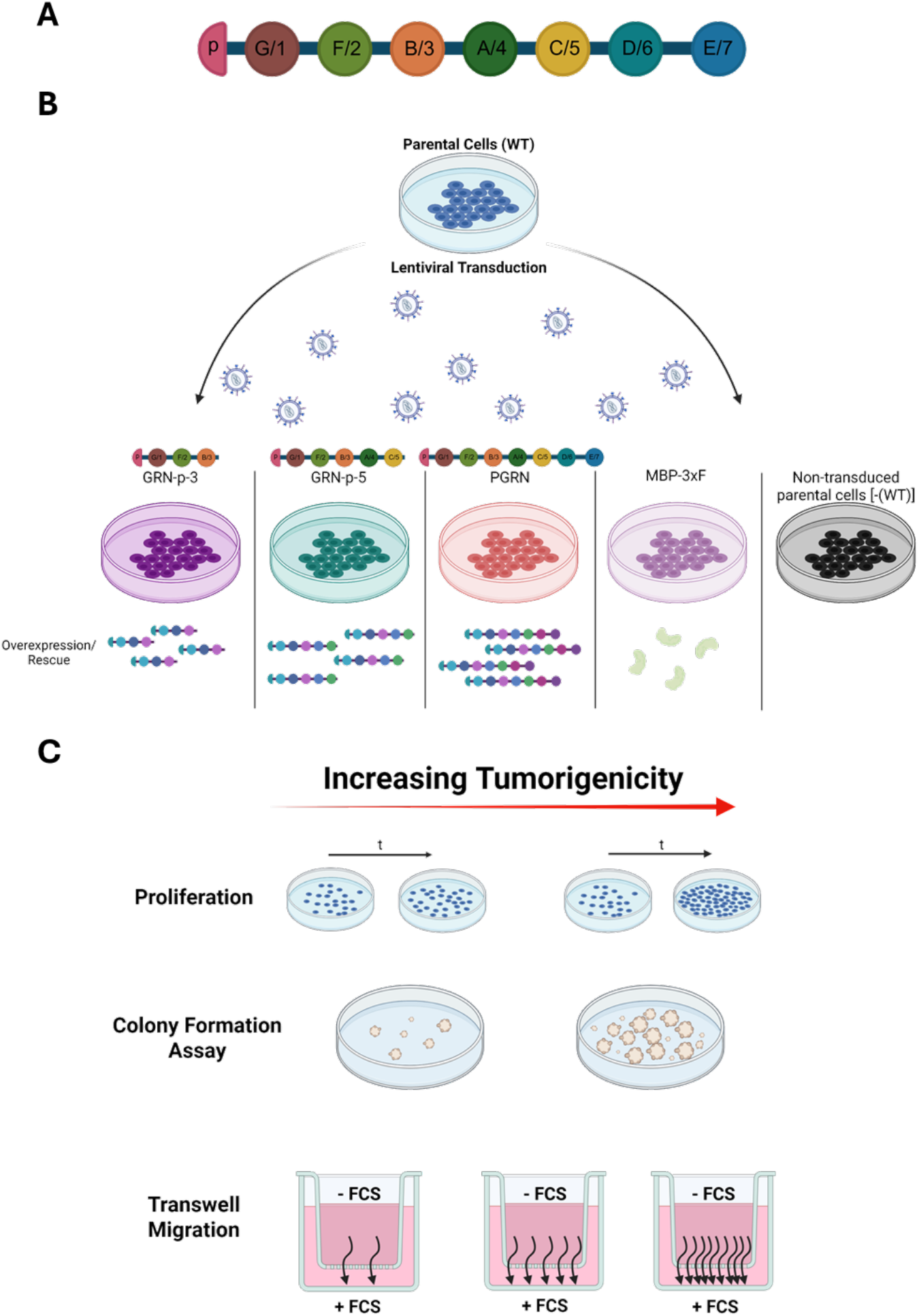
Rationale and planned experiments. (A) Schematic illustration of the PGRN protein with the seven and a half GRN motifs depicted. (B) Scheme displaying the lentiviral overexpression constructs coding for the indicated portion of PGRN or for 3xFLAG-tagged maltose-binding protein (MBP 3xF) as a control. (C) Graphical illustration of the experiments used to assess the tumorigenicity of cultured cell lines: measuring cell proliferation over time, the ability to form colonies under low serum conditions or in soft agar (anchorage-independent growth), and transwell migration assays.

In the context of cancer, most of the functional data stems from treating the cells with full-length PGRN or knocking down PGRN. Targeting PGRN was proposed as a potential therapeutic approach, because its downregulation has been shown to increase apoptotic markers and the sensitivity to chemotherapeutics, while the upregulation leads to increased levels of cancer stem cell markers ^13–16^. Targeting PGRN might be an interesting approach in cell-based assays; its feasibility in patients, however, depends on the precision in targeting cancers and the temporal availability, as systematic downregulation could lead to severe neurological effects and increased inflammation ^2,6,7^.

While PGRN might be a difficult target for therapeutic options, it shows great promise as a biomarker. PGRN blood levels positively correlate with aggressiveness and higher chances of metastasis for a broad range of cancers, including breast cancer ^17^, papillary thyroid carcinoma ^13,18^, and prostate cancer ^19^. PGRN is detected in the urine, and PGRN levels in the cerebrospinal fluid are a well-characterized indicator for metastasis in the central nervous system (CNS) ^20^.

Here, we aimed to gain functional data, dissecting which GRNs drive cancer development or whether full-length PGRN is required for increased proliferation, migration, and metastasis formation. As the example of Atsttrin showed, the effect of PGRN and/or individual GRNs might be mediated by a combination of different GRNs. We created lentiviral vectors to overexpress C-terminally truncated versions of PGRN that lack the last two (GRN-p-5) or the last four GRNs (GRN-p-3; Figure 1B). We transduced *wild-type* and *GRN*-KO HepG2 (hepatoma cell line) and U87 (glioblastoma cell line) cells with these different truncated PGRN-expressing lentiviruses and assessed the performance of the cells in cell-based tumorigenicity assays, such as colony formation, migration, and proliferation, both in normal cell culture media and in media with reduced serum concentration (Figure 1C).

Elevated PGRN has been described in both hepatocellular carcinoma patients and glioblastoma patients ^21,22^. HepG2 cells treated with shRNA targeting the PGRN mRNA were shown to grow more slowly and perform worse in cell-based tumorigenicity assays ^22^. U87 cells were shown to express more PGRN than glial and astrocyte cell lines, and the glioblastoma cell line S1R1 showed a PGRN-dependent increase in tumorigenicity in cell culture experiments ^21^. We therefore wanted to test whether this also holds for U87 cells.

While overexpression of full-length PGRN caused an increase in colony formation in HepG2 cells, we did not observe a similar effect in the U87 cells. Overexpression of the truncated PGRN constructs GRN-p-5 and GRN-p-3 did not significantly increase in any of the performed assays. Contrary to the previously published data described above, we did not observe an increase in the proliferation of HepG2 cells upon progranulin overexpression. Moreover, the proliferation of U87 cells was not affected by increasing or decreasing PGRN levels.

## Results

### Lentiviral transduction leads to stable overexpression of PGRN constructs

HepG2 and U87 *wild-type* cells were transduced with lentiviral expression constructs depicted in Figure 1, and the expression of the various constructs in HepG2 cells was confirmed by western blotting (Figure 2A). While the cells transduced with the full-length PGRN and the GRN-p-5 encoding lentiviruses showed clear overexpression, the signal from the GRN-p-3 construct was weaker (Figure 2A). This weaker signal might represent reduced expression of GRN-p-3 and/or be due to the fewer available epitopes for the used polyclonal antibody to bind to on the shorter GRN-p-3 constructs.

**Figure 2:**
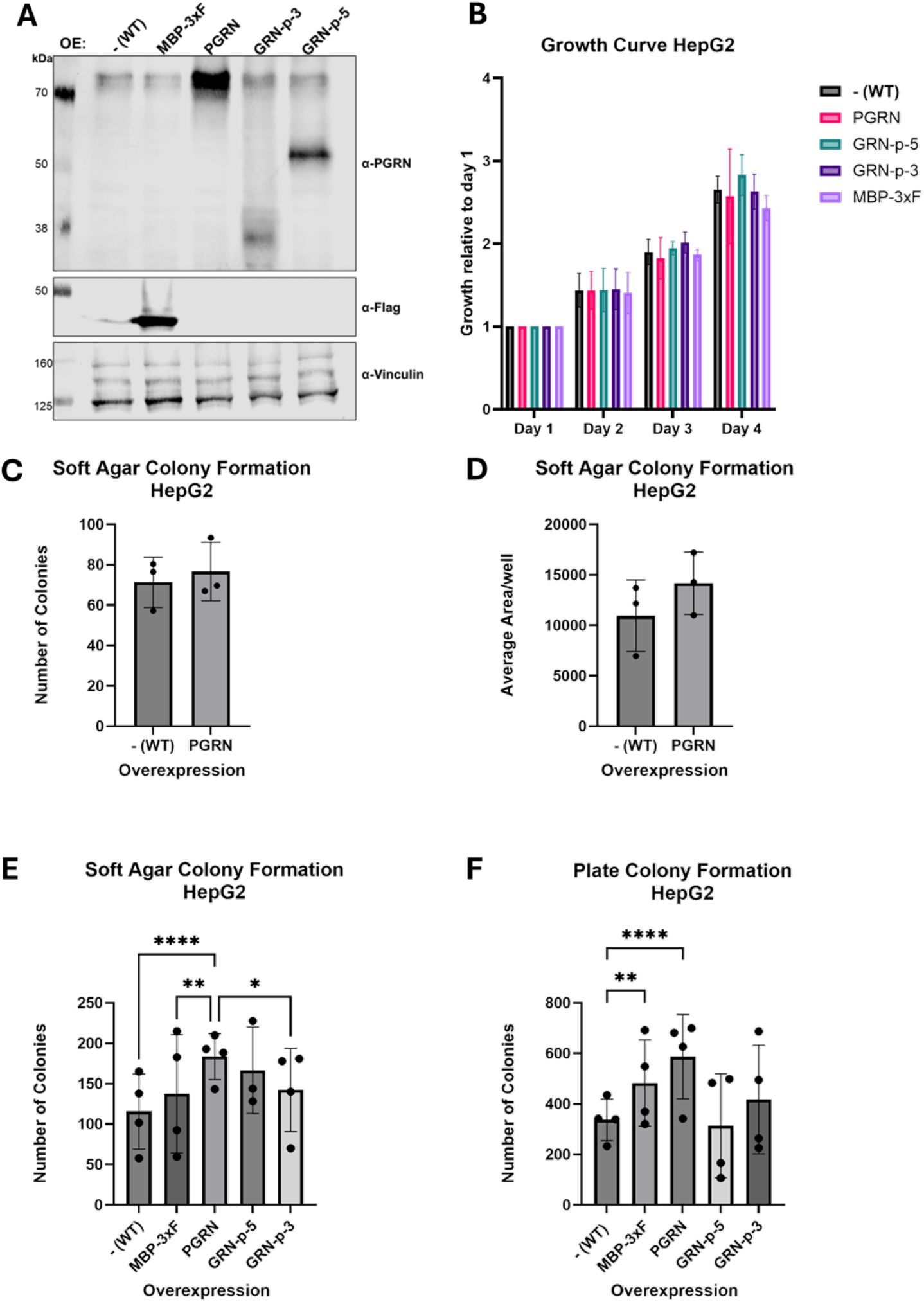
Progranulin overexpression in HepG2 cells. (A) Western Blot confirming overexpression (OE) of MBP-3xF and progranulin constructs in HepG2 cells. The non-transduced parental cells are denoted –(WT). The effects of the different overexpression constructs were then assessed in proliferation assays (B) and in soft agar colony formation assays, where the number of colonies (C, E) and the colony size (D) were quantified. (F) Number of colonies in plate colony formation assays of HepG2 cells, where cells need to grow in 1% FCS. Results in E and F were subjected to ordinary two-way ANOVA followed by Tukey’s multiple comparisons test with significance levels (^*^≤0.01, ^**^≤0.001, ^****^≤0.0001). Data are represented as mean +/-standard deviation (SD). Each data point was calculated as the mean of 6 technical replicates. B, C, D, and E were created using GraphPad Prism.

### Full-length PGRN overexpression increases the growth of HepG2 cells in soft-agar and plate colony formation assays

After confirmation of expression, we first analysed the effect of the lentiviral PGRN constructs on cell proliferation and the colony-forming ability in HepG2 cells. To assess proliferation, we used the colorimetric CellTiter 96^®^ AQueous One dye in a time course experiment over 4 days. The amount of metabolized dye was detected by mseasuring absorbance at 490 nm after one hour of incubation and serves as a proxy for the number of cells present in each well. The reference measurement (Day 1) was conducted one day after cell seeding (Day 0). Proliferation was then quantified as an increase in absorbance compared to the value on day 1. None of the PGRN or GRNs overexpressing HepG2 cells showed a statistically significant increase in proliferation compared to the non-transduced parental [-(WT)] or the MBP-3xF overexpressing HepG2 cells (Figure 2B). Prolongation of the measuring time to 7 days did not change the differences between the cell lines (data not shown).

Next, we tested the ability of the HepG2 cells to form colonies under low serum conditions (1% FCS) in the plate colony-formation assay and in the soft agar colony-formation assay, where cells must grow in an anchorage-independent manner, embedded in a mix of culturing media and agarose. Compared to the WT cells, the PGRN overexpressing cells showed a small but statistically not significant increase in the number of colonies formed and in the overall size of the colonies in the soft agar assay (Figure 2C-D). Repetition and further comparison of PGRN with the control cells [-(WT) and MBP-3xF] confirmed this trend and showed a statistically significantly increased number of colonies of the PGRN cells in soft agar (Figure 2E). While both GRN-p-5 and GRN-p-3 showed a small increase, too, the effect was not statistically significant (Figure 2E).

In the plate colony formation assay, growth of PGRN cells was significantly increased, compared to the parental cells [-(WT)] and MBP-3xF. The variability between biological replicates in the GRN-p-5 and GRN-p-3 was high, and on average, no significant difference in the number of colonies in WT and MBP-3xF was observed (Figure 2F).

Migration has previously been reported to be affected by progranulin levels in HepG2 cells ^22^. We assessed migration both in transwell migration assays and in scratch assays. In the transwell migration assay, we did not observe any directed migration in general for HepG2 (data not shown), and also in the scratch assay, we only observed minimal migration for the different cell lines (Supplementary Figure S1). Overall, it is questionable whether the closure of the scratch was due to proliferation or actual migration of HepG2 cells.

### PGRN overexpression in U87 does not increase proliferation

After confirming the overexpression of our full-length PGRN and oligo-GRNs constructs in U87 cells (Figure 3A), we measured cell proliferation by performing growth curves as with the HepG2 cells. In the U87 cells, overexpression of the different constructs did not alter cell proliferation compared to the non-transduced parental cells [-(WT)] (Figure 3B).

**Figure 3:**
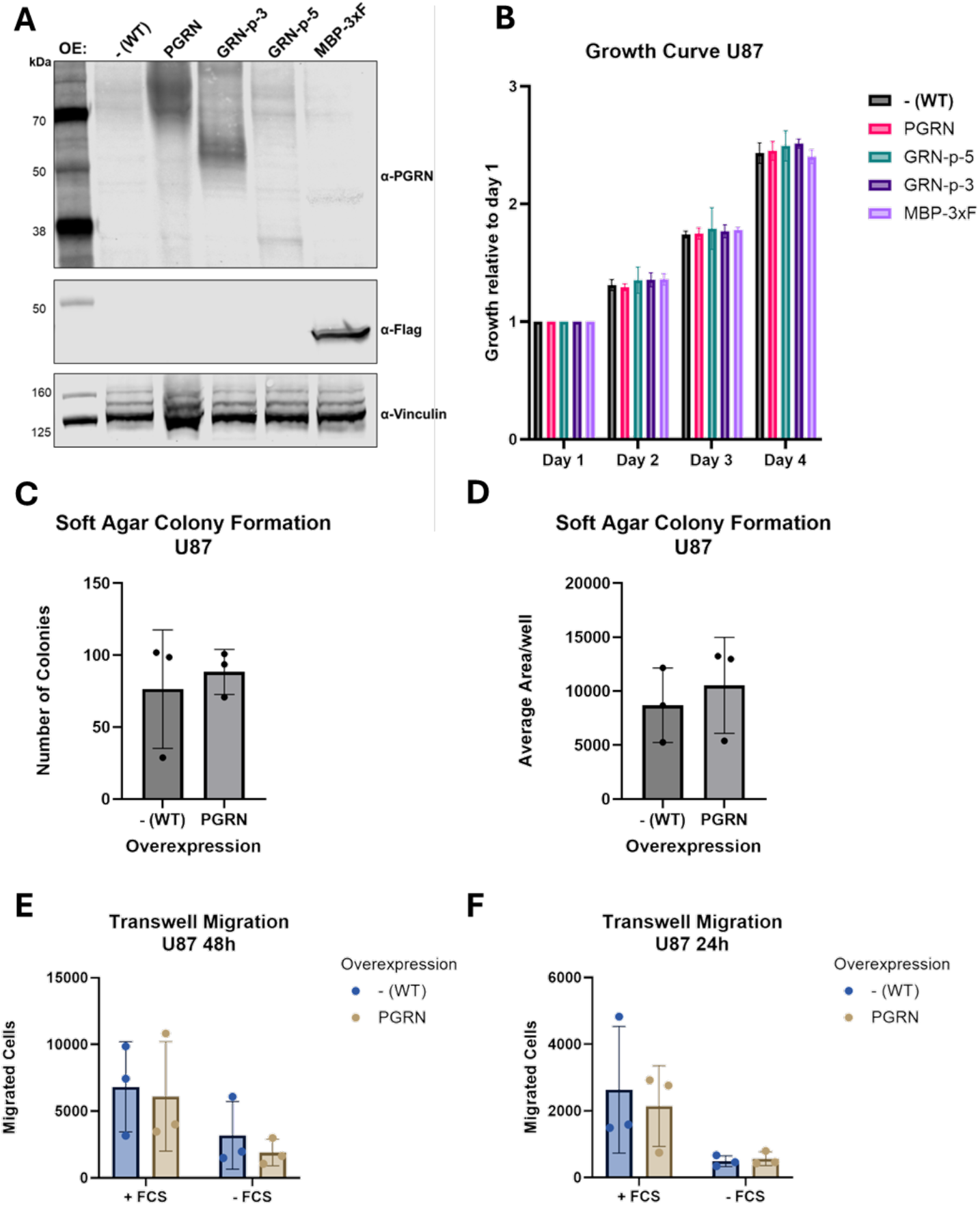
Progranulin overexpression in U87 cells. (A) Western Blot confirming overexpression of MBP-3xF and the full-length and truncated PGRN constructs in U87 cells. The effect of the various overexpression constructs was assessed in proliferation assays (B). Soft agar colony formation assays, quantifying both, the number of colonies (C) and the colony size (D), as well as in transwell migration assays (E, F), where cell migration through the pores of a membrane was measured, either from serum free medium towards medium with 10% FCS (+FCS) to assess chemoattractant-mediated directional migration or random migration towards serum-free medium (−FCS), were conducted with full-length progranulin overexpressing (PGRN) and non-transduced parental cells [-(WT)]. B, C, D, E, and F were created with GraphPad Prism.

Comparing anchorage-independent growth in the soft agar colony-formation assay between PGRN and parental U87 cells showed a slight increase in both the number of colonies and the mean size of these colonies. However, this increase was not statistically significant (Figure 3C-D).

U87 cells generally grow fast and have a high migratory potential. Therefore, the plate colony formation assay could not be performed, as the cells did not form colonies and migrated throughout the plate.

To quantify the migratory potential of these cells, we next performed transwell-migration assays. Cells were therefore seeded in the upper part of the transwell inlay, in media devoid of serum, while the lower part of the well contained complete media containing 10% FCS (Figure 3E-F; +FCS) or media without serum (−FCS) as a control for random, non-directional migration. While after 48 hours, we observed an extensive amount of undirected cell migration even in the -FCS control condition, after 24 hours, there was a clear distinction between directed (+FCS) and undirected migration (−FCS). However, in all conditions, there was no significant difference in migration activity between the parental and the PGRN overexpressing U87 cells (Figure 3E-F).

### siRNA mediated knockdown of PGRN in U87

Elevated PGRN levels have been reported to lead to increased tumorigenicity, whereas decreased PGRN has been shown to reduce proliferation, cell migration, and anchorage-independent growth ^21–23^. To knock down endogenous PGRN, we designed siRNAs targeting the 3’ untranslated region (UTR) of the PGRN mRNA. Following siRNA transfections, the knockdown efficiency was assessed using RT-qPCR (Figure 4A) and western blotting (Figure 4D). The knockdown efficiencies achieved with siGRN1, siGRN3, or a mix of both were similar to another frequently used siRNA in our lab, which targets the protein upstream frameshift 1 (UPF1) and which was used here as a specificity control (Figure 4B). PGRN mRNA was depleted 10-fold 48 hours after the second transfection, and the knockdown did not affect the overexpressed PGRN, which lacks this portion of the 3’ UTR (Figure 4C-D).

**Figure 4:**
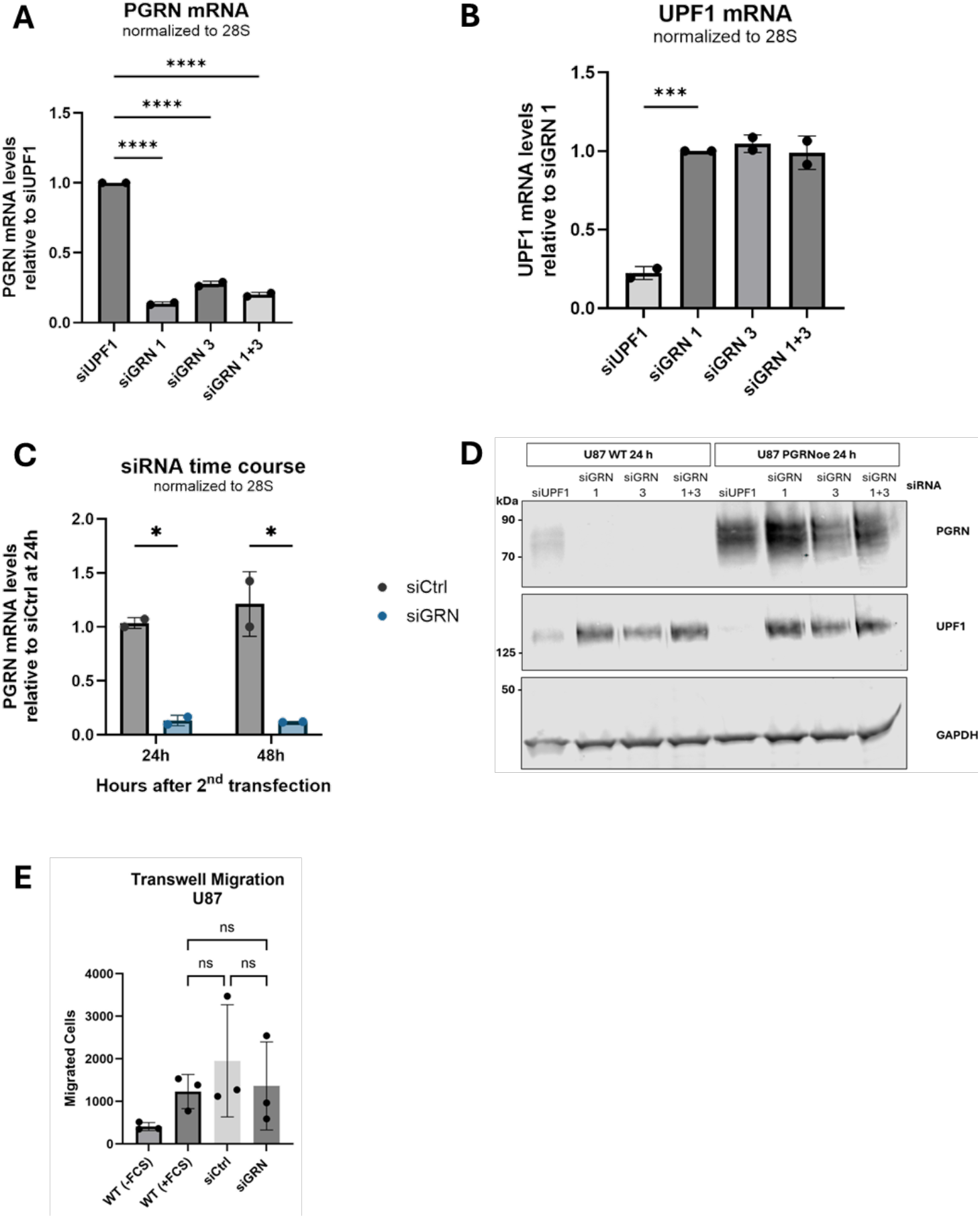
siRNA mediated knockdown of progranulin. (A) PGRN mRNA levels 24 h after the 2nd siRNA transfection, relative to siUPF1 treated control cells and normalized to 28S rRNA. (B) UPF1 mRNA levels 24 h after the 2nd siRNA, relative to siGRN 1 treated U87 cells and normalized to 28S rRNA. (C) PGRN mRNA levels in non-targeting control (siCtrl) and GRN siRNAs (siGRN 1+3) treated cells 24 h and 48 h after the 2nd siRNA transfection, relative to siCtrl treated cells after 24 h. (D) Western blot of U87 WT and PGRN overexpressing cells confirming knockdown of endogenous GRN 24 h after the 2nd siRNA transfection. U87 cells were treated with siRNAs against UPF1 (siUPF1, positive control) or endogenous GRN (siGRN 1, 3, 1+3). GAPDH served as a loading control. (E) Transwell migration assay showing the number of migrated U87 WT or siRNA (siGRN 1+3, siCtrl) treated cells towards medium with FCS chemoattractant (+FCS) and random migration of U87 WT cells towards serum-free medium (−FCS). Statistical significance was determined by ordinary one-way ANOVA followed by Dunnett’s multiple comparisons test (A, B) or repeated measures one-way ANOVA followed by Tukey’s multiple comparisons test (E). Results in C were subjected to ordinary two-way ANOVA and Šídák’s multiple comparisons test. ns (non-significant), ^*, **, ***^, and ^****^ indicate p-values of >0.01, ≤0.01, ≤0.001, ≤0.001 and ≤0.00001, respectively. A, B, C, and E were created using GraphPad Prism.

We tested whether the knockdown of endogenous PGRN in U87 cells affected their migratory potential in the transwell migration assay. As a control, we transfected the cells with a non-targeting control siRNA. However, during the 24 hours of the migration assay, the U87 cells were not affected by the absence of PGRN (Figure 4E).

### Granulin knockout by CRISPR/Cas9 affects growth independently of progranulin

To test the effect of a complete and long-term PGRN depletion, we generated clonal *GRN* knockout cell lines from HepG2 and U87 cells using CRISPR/Cas9 genome editing with gRNAs targeting exons 4 and 9. After transfection and selection, clones were screened for low PGRN mRNA levels by RT-qPCR, and *GRN* knockouts were then confirmed by western blotting (Figure 5A). All clonal cell lines originate from single cells, ensuring homogenous genotypes. We then transduced the *GRN*-KO cells with the previously described lentiviral vectors and confirmed rescue and overexpression of the various PGRN constructs by western blotting (Figure 5B).

**Figure 5:**
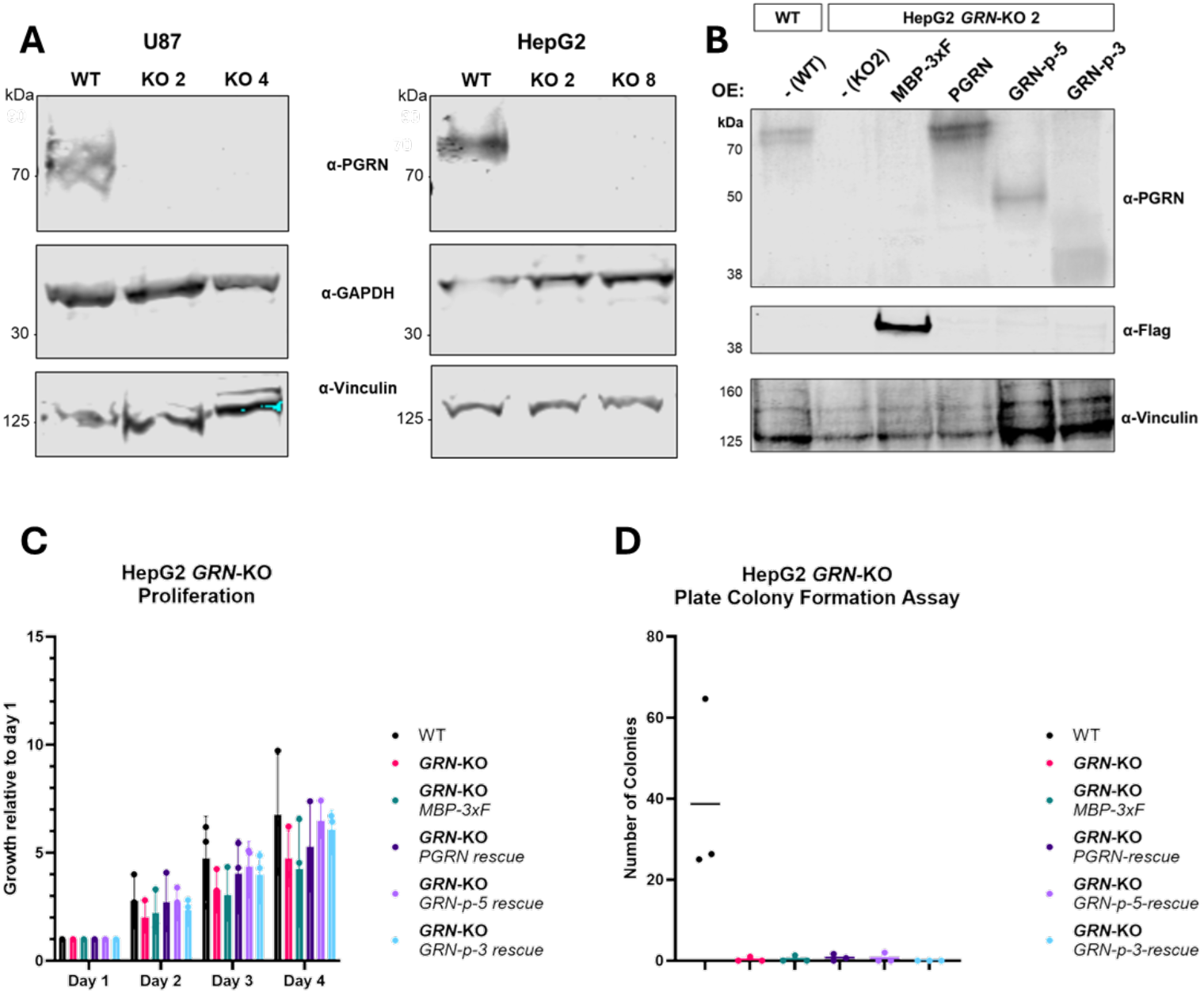
CRISPR/Cas9 knockout of progranulin. (A) Western blots confirming the homozygous knockout (KO) of the GRN gene in two clones of U87 and HepG2. GAPDH and vinculin serve as loading controls. (B) Western blot confirming GRN-KO and rescue in HepG2 cells (clone 2). Expression of PGRN, GRN-p-5, and GRN-p-3, as well as the MBP-3xF control in the respective HepG2 cell lines. Vinculin serves as loading control. (C) Proliferation of HepG2 GRN KO and rescue cell lines and (D) plate colony-formation assay. C and D were created with GraphPad Prism.

Next, we tested the proliferation of HepG2 parental (WT), *GRN*-KO, and rescue cell lines. The HepG2 *GRN*-KO cells showed reduced growth (Figure 5C) compared to parental cells. However, the growth of the control cells transduced with MBP-3xF was similarly reduced. The reduced proliferation was rescued to some extent by all PGRN and oligo-GRN expression constructs. However, the variation among the biological replicates was considerable, rendering the effects statistically non-significant (Figure 5C).

Overall, the HepG2 knockout cells were more sensitive to stress and showed worse survival during the lentiviral transduction and the following selection. This sensitivity was also observed in the plate colony-formation assay, where all cell lines that underwent genome editing did not survive the low FCS concentration (Figure 5D). Serum concentrations below 5% lead to reduced growth and increased cell death, independently of the progranulin expression, hinting at effects originating from the genome editing, as the effect was observed in both *GRN*-KO clones 2 (Figure 5C-D) and 8 (data not shown).

While the *GRN*-KO HepG2 cells suffered and showed reduced growth, we observed a PGRN-independent proliferation increase in U87 cells that underwent the CRISPR/Cas9 genome editing (Supplementary Figure S2A-B).

## Discussion

In this study, we confirmed the PGRN-mediated increase in both anchorage-independent growth and colony formation under low serum conditions (1% FCS) in HepG2 cells, as previously reported by Liu and colleagues ^22^. However, we were not able to reproduce other phenotypes they reported in their study. The most striking difference concerns the migration assays, in which our HepG2 cells exhibited very low migration activity. In the scratch assay (also known as wound healing assay), we did not see a PGRN-dependent effect on migration (Supplementary Figure S1). We suspect that the previously reported faster closure of the scratches in PGRN overexpressing or progranulin-treated HepG2 cells in the scratch assay ^22^ might not result from differences in migration but rather could be caused by increased cell proliferation. This explanation is consistent with the reported elevated cell proliferation in that study. Additionally, to exclude contaminated cultures, we confirmed that our cells are indeed HepG2 cells, using the cell line authentication service from Microsynth AG, which relies on short tandem repeat (STR) analysis.

While for HepG2 cells, phenotypes have been shown for both PGRN depletion and overexpression, we wanted to expand the cellular toolbox. Bandey and colleagues ^21^ showed a dependence of the glioblastoma cell line S1R1 on PGRN, both with regard to the sensitivity to chemotherapeutics and in the overall fitness. For U87 cells, they reported elevated PGRN levels, compared with non-tumorigenic glia cells, but did not perform further characterization of U87 cells. We observed neither an effect of PGRN depletion nor overexpression on overall cell survival, growth, or migratory potential.

This highlights that PGRN can have a pro-tumorigenic effect, but that it is not necessarily required to promote growth and migration in all cancer cell lines. While most cancer-related publications on PGRN report an effect of PGRN, studies observing no effect are more likely not to have been published, which overall generates a skewed picture of PGRN’s involvement in cancer ^24^.

Our attempt to investigate the roles of the various GRN motifs in promoting proliferation and growth in cancer was not entirely successful. While we detected trends of small increases in cell growth when full-length PGRN was overexpressed, we were not able to detect potentially smaller effects resulting from overexpression of the C-terminally truncated constructs GRN-p-5 and GRN-p-3. Some interactions between PGRN and various proteins have been mapped to different granulins or combinations of granulins: For example, Sortilin has been shown to bind to the C-terminus of granulin E ^25^, Prosaposin to granulins D and E ^11^, and TNFR to Atsttrin, which consists of granulins F, A, and C plus some linkers ^3^. Thus, it would be interesting to see whether some GRNs are dispensable for increasing tumorigenicity. This would potentially allow ruling out the involvement of some PGRN interactors as mediators for its tumorigenic potential. While the expression of different deletion mutations of progranulin either in the context of overexpression or rescue remains interesting, it might be necessary to investigate this in other cell lines, where PGRN appears to be essential for tumorigenicity, such as S1R1 ^21^ or Hep3B ^22^.

Many pieces in the puzzle around the biology of PGRN and individual GRNs are still missing. The pleiotropic nature of PGRN makes the distinction between different functions of individual GRNs or PGRN in different pathways challenging. PGRN knockdown was proposed as a potential cancer therapeutic to sensitize the cells to conventional anti-cancer therapies, whereas on the other side of the spectrum, the exogenous supplementation of PGRN was suggested as a therapy for Frontotemporal Dementia ^26–29^. Therefore, more fundamental research to elucidate PGRN functions is needed to understand and address potential side-effects of the proposed therapeutic approaches and to develop a clear safety profile.

## Competing interests

The authors declare no competing interests.

## Author contributions

AH, M-DR, and OM conceived the project; AH and LT planned, executed, and analysed the experiments. AH, LT, OM, and M-DR interpreted the results. AH composed all figures. AH, M-DR, and OM wrote the manuscript.

## Acknowledgments

We are grateful to Dr. Sofia Nasif for her input on protocols and for helping to troubleshoot, to Prof. Dr. Paola Luciani and Prof. Dr. Eric Vassella for providing the HepG2 hepatoma and the U87 glioblastoma cell lines, respectively, to Dr. Niamh O’Brien for support and guidance on CRISPR/Cas9 genome editing, to Tomas Solomon for advice on generating lentiviral particles, and to Karin Schranz for her help with cell culture experiments.

## Funding

This work was funded by the National Center of Competence in Research (NCCR) on RNA & Disease, funded by the Swiss National Science Foundation (SNSF; grant 51NF40-141735), by the SNSF grant 310030-204161 to O.M., and by the canton of Bern (University intramural funding to O.M.). This research was also made possible through the support of the UK Dementia Research Institute through UK DRI Ltd (award number UK DRI-6005), funded by the UK Medical Research Council, Alzheimer’s Society and Alzheimer’s Research UK) and (award number UK DRI-6204), principally funded by the UK Medical Research Council) to M-DR.

## Materials and Methods

### Cell Culture

Cells were cultured in DMEM/F-12 (Gibco, 32500) supplemented with 1x Penicillin/Streptomycin (Gibco, 15070063) and 10% FCS (BioConept, 2-01F10-I), further referred to as DMEM (+/+). To passage, cells were washed with D-PBS (Gibco, 14190-094) and detached using Trypsin-EDTA PBS (T/E) (BioConcept, 5-51F00-H).

For *Soft Agar Colony-Formation Assays*, a bottom layer of 1.5 mL 1x DMEM (+/+) containing 0.5% agarose (Promega, V3125) was casted in each well of a 6-well plate. After solidifying, 10’000 cells were seeded in the upper layer of 1.5 mL of DMEM (+/+) containing 0.3 % agarose. Cells were then cultured for three weeks, and 0.5 mL media was added and changed twice per week. After three weeks, the media was removed, each well was washed once with D-PBS and then fixed and stained with 0.5 mL of 0.01% crystal violet in 20% methanol for one hour. The agar was then de-stained with water, until only the colonies remained stained. Wells were imaged using the GelDoc from Vilber, and colonies were analysed using Fiji ^30^.

For *Plate Colony-Formation Assays*, 10’000 cells were seeded in DMEM/ F-12 supplemented with 1x P/S but only 1% FCS. The cells were cultured for two weeks and then fixed, stained, and analysed as explained for the soft agar colony-formation assay.

For *Transwell Migration Assays*, 50’000 cells per transwell inlay were seeded in 0.5mL DMEM/F-12 devoid of FCS but containing 1x P/S (DMEM (+/-). In the lower compartment, either DMEM (+/+) or DMEM (+/-) was added and the cells were incubated for 24-48 hours. The transwell was then washed and the cells in the upper part of the transwell were removed with a cotton swab. The cells on the lower part of the membrane were fixed in 4% paraformaldehyde for 20 minutes and stained with DAPI. After staining, the membrane was washed with PBS, cut out using a scalpel, and further placed on a microscopy glass slide. Mounting media was pipetted on top of every membrane, and a cover slip was placed on top, which was further sealed with nail polish. Images were taken using a fluorescent microscope, and numbers of migrated cells were determined with Fiji ^30^.

*Cell proliferation* was assessed using the colorimetric CellTiter dye from Promega (also referred to as cell viability assay), following the manufacturer’s protocol. In short, 10’000 HepG2 cells, or 3’000 cells in the case of U87 and H4, were seeded in 0.1 mL DMEM (+/+) per well of a flat-bottom 96-well plate. 24 hours after seeding, the first measurement was taken. All further measurements were then normalized to the first measurement to account for seeding differences. For analysis, the media was changed at least 2 hours before incubation with the dye. 20 μL of the CellTiter dye was then added to each well, and the cells were incubated for one hour. Thereafter, the absorbance was measured at 490 nm in the Tecan Infinite M1000 plate reader.

### Western Blot

For Western Blots (WB), cells were lysed in RIPA buffer (10 mM Tris-Cl (pH 8), 1 mM EDTA, 1% Triton X-100, 0.5% sodium deoxycholate, 0.1% SDS, 150 mM NaCl), supplemented with 1x protease inhibitor (ABSOURCE, B14002), for 30 minutes on ice. Protein concentration was measured using the Direct Detect equipment (Millipore), and equal amounts of protein were incubated at 70 °C in 1x SDS running buffer for 10 minutes. For PGRN detection, samples were run on a self-cast 8% PAA gel, otherwise mPAGE 4-12% Bis-Tris gels (Millipore) were used. For PGRN detection, it is important not to use reducing agents, such as TCEP or DTT, in the buffers. Primary and secondary antibodies, along with the dilutions at which they were used, are shown in Table 1. Membranes were incubated with primary anti-bodies overnight at 4 °C, washed three times for 5 minutes with TBS-t, before being incubated with secondary antibodies at room temperature for one hour and thereafter washed again three times with TBS-t. All antibodies were diluted in TBS-t containing 5% milk. Membranes were scanned using the odyssey scanner from Licor.

**Table 1:** Antibodies for Western Blot.

| **Primary Antibodies** |  |  |  |  |
| --- | --- | --- | --- | --- |
| **Article** | **Dilution** | **Clonality** | **Catalog Nr.** | **Supplier** |
| Goat anti-PGRN | 1:250-500 | Polyclonal | AF2420 | R&D Systems |
| Mouse anti-Flag | 1:1000 | Monoclonal | F1804 | Sigma-Aldrich |
| Mouse anti-Vinculin | 1:2000 | Monoclonal | Sc-73614 | Santa Cruz |
| Mouse anti-GAPDH | 1:10000 | Monoclonal | Sc-47724 | Santa Cruz |
| **Secondary Antibodies** |  |  |  |  |
| **Article** | **Dilution** | **Fluorescence** | **Catalog Nr.** | **Supplier** |
| Donkey anti-Goat IgG | 1:10000 | IRDye® 800CW | 926-32214 | Licor |
| Donkey anti-Mouse IgG | 1:10000 | IRDye® 680RD | 926-68072 | Licor |

### Lentiviral Transduction

#### Lentivirus production

The parental plasmid pLVX-EF1a-TS-delta56-110FUS-IRES-PuroR, generated from pLVX-EF1a-IRES-PuroR (Clontech, Cat. Nr. 631988) and described in Humphrey and colleagues ^31^, was digested with SpeI and BamHI, and the cut vector was run on a gel and afterwards cleaned up. The full-length PGRN insert was amplified from cDNA by PCR, using Ex Premier DNA Polymerase (TaKaRa RR370). The MBP-3xF was amplified from previously used plasmids in the lab ^32^. Both inserts were further ligated in the digested vector using the In-Fusion Cloning kit by Takara, following the manufacturer’s instructions. The respective primer sequences are shown in Table 2.

**Table 2:**
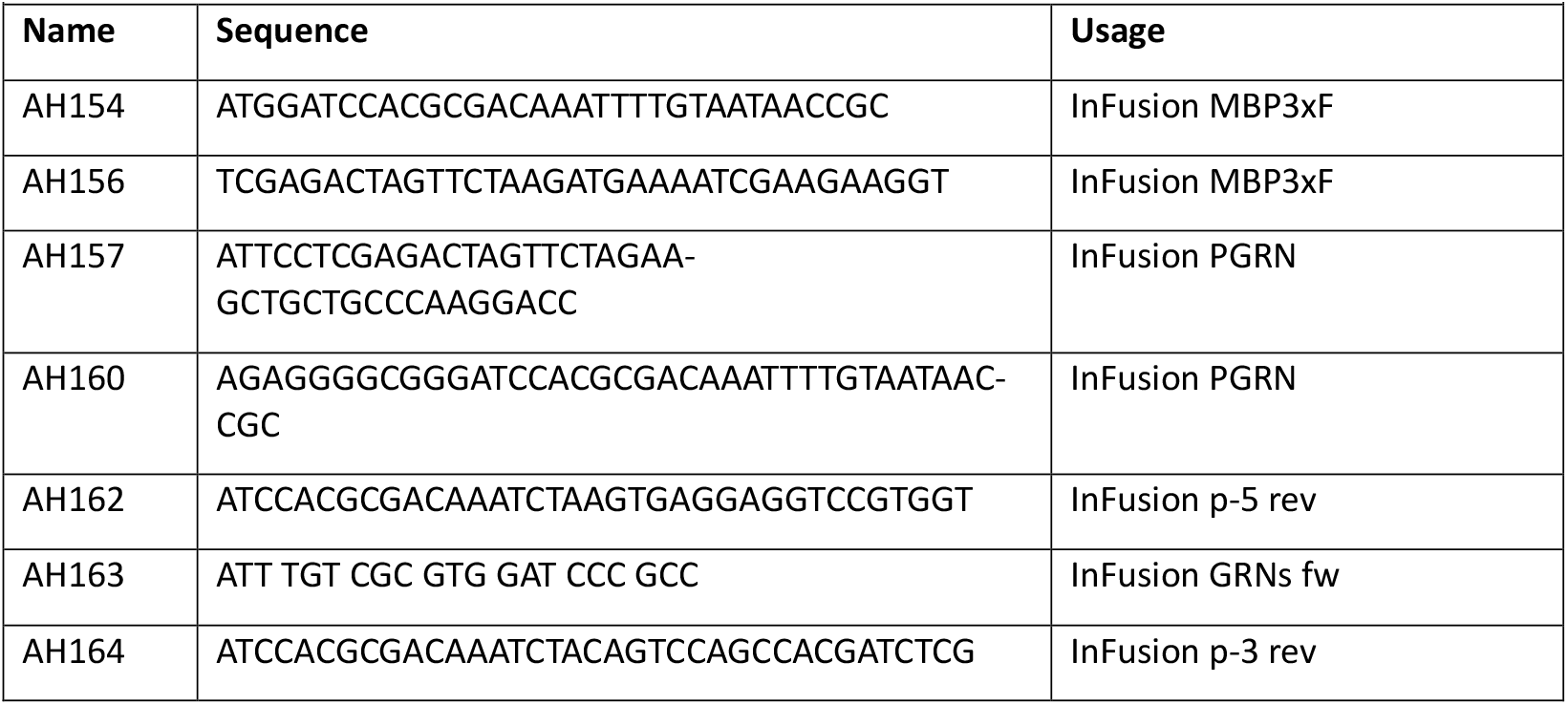
PCR Primers for pLVX InFusion Cloning.

For the GRN-p-5 and GRN-p-3 expression vectors, which lack the last two and last four GRNs, respectively, we amplified the pLVX-EF1a-PGRNoe backbone by PCR, leaving out the unwanted GRNs, using AH163 as forward primer in both cases, and AH162 and AH164 as reverse primers for the GRN-p-5 and GRN-p-3, respectively (Table 2). The resulting PCR product was ligated using the InFusion Kit (Takara). All Lentiviral plasmids were transformed into Stbl3 bacteria (Invitrogen).

Lentiviral particles were produced using the Trans-Lentiviral Packaging kit (Dharmacon Horizon, TLP5919). In short, HEK293T cells in a T75 flask (TPP, 90076) were transfected at a confluency of ∼70% with 15 μL Trans-Lentiviral Packaging Mix,11 μg Lentiviral Vector, and 90 μL of Lipofectamine 2000 Invitrogen, 11668019). The transfection complexes were incubated in OptiMem (Gibco, 51985-034) before transfection. Media was changed 24 h after transfection to 15 mL of fresh DMEM/F-12 (+/+). On the following days, 48 h and 72 h, the media was harvested and filtered through CA syringe filters (Avantor vwr, 514-1271). 10 mL (ratio 1:4) of Lenti-X-Concentrator (Takara, 631232) was added to the cold media and stored at 4 °C for 30 min to 3 days, before being centrifuged at 4 °C for 45 min at 1500 rcf.

The supernatant was decanted, and the pellet was resuspended in 2 mL cold PBS. The viral particles were transferred to cryotubes. Virus titres were determined using the Lenti-X GoStix Plus lateral flow tests (Takara, 631280) and the cryotubes were snap frozen in liquid nitrogen, before being stored at - 80 °C for later use.

#### Lentiviral transduction

For transduction, 2-4 ^*^ 10^5^ cells were seeded in a 6-well plate. The following day, the media was replaced with fresh media containing a final concentration of 5 μg/mL polybrene. Lentiviruses were diluted to a GoStix Value (GV) of 250-500 ng/mL virus particle protein p24. The volume corresponding to 25 ng p24 was added to each well of a 6-well plate. 24 hours after addition of the virus, media was changed to fresh media, and 48 hours after transduction selection was started with Puromycin (1 μg/mL for HepG2 and 0.8 μg/mL for U87). When reaching confluency, cells were split and further cultured in selection media until the no-transduction control showed no more surviving cells.

### CRISPR/Cas 9 GRN knockouts

For the CRISPR/Cas9 genome editing, we used a 2-plasmid approach consisting of one plasmid coding for both the Cas9 and the sgRNA (described in Reber et al. ^33^) and a EF1a-pRR-PuroR plasmid ^34^, which results in transient resistance of the cells to puromycin when the plasmid is cut by the Cas9 and undergoes homologous recombination. The EF1a-pRR-PuroR plasmid was cloned by annealing two oligos (Table 3), containing the gRNA binding site, followed by restriction digestion and ligation into the pRR-PuroR plasmid using SalI and SpeI, as described elsewhere ^35^.

**Table 3:**
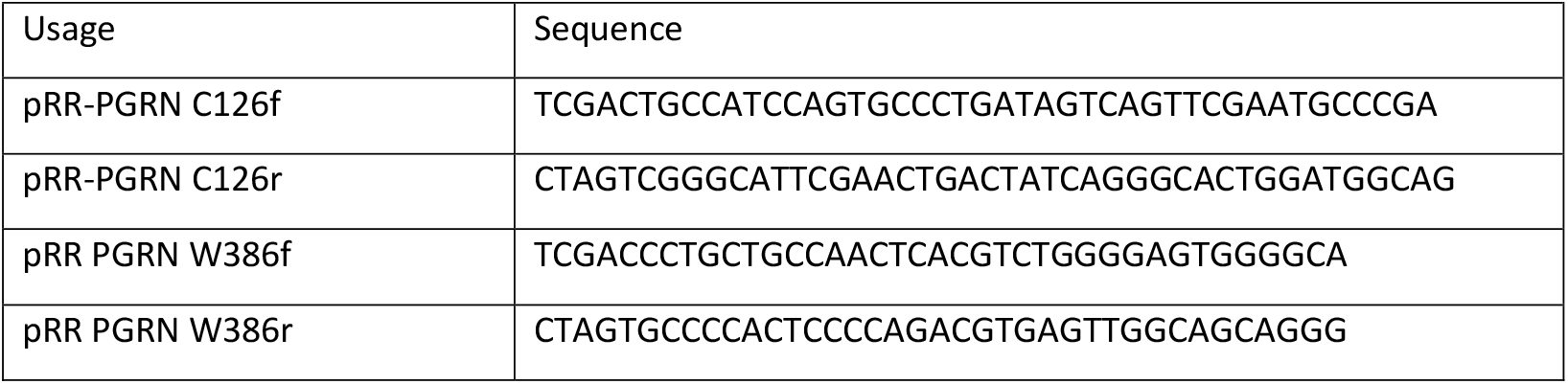
Oligos for pRR-PuroR cloning.

We used two different guideRNAs (gRNAs), targeting *GRN* p.W386 (GCCAACTCACGTCTGGGGAGTGG) and p.C126 (GGCATTCGAACTGACTATCAGGG). 48 hours after transfection, the cells were transferred to a 15 cm dish and selected with 0.5 µg/mL puromycin. After selection, cells were grown until colonies reached > 20 cells, at which point the colonies were picked from the plate using a P1000 pipette, expanded, and ultimately cryopreserved. PGRN mRNA levels were assessed by RT-qPCR using the Brilliant III UF SYBR Green qPCR Master Mix (Agilent, 600882) and the qPCR primers indicated in Table 4, followed by further assessment of truncated PGRN levels by western blotting.

**Table 4:**
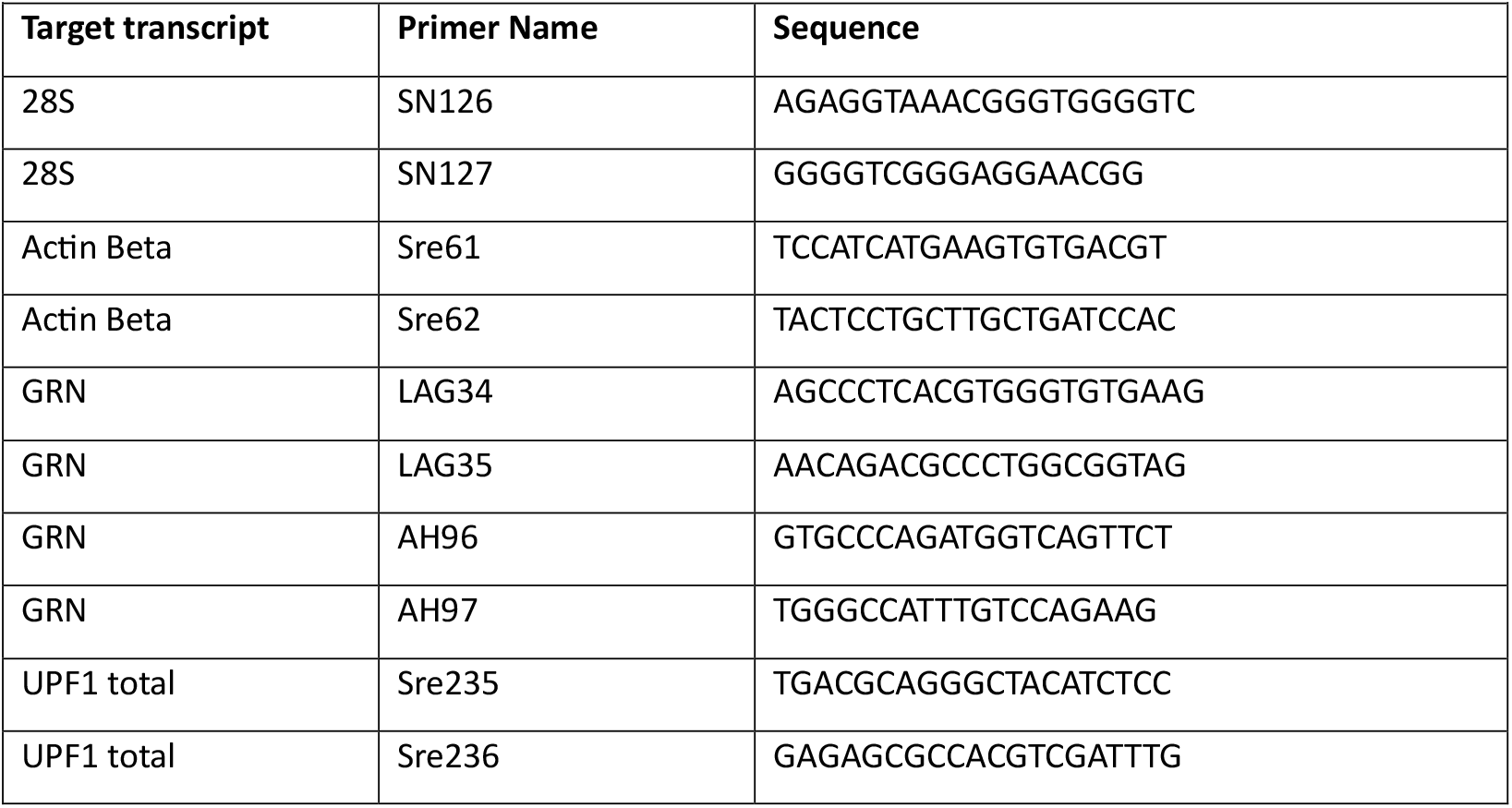
qPCR primers.

### siRNA-mediated knockdowns

For the PGRN knockdown with siRNAs, 100’000 cells were seeded per well of a 6-well plate on day 0. On day 3, 1.5 mL of fresh media was added to the cells and the cells were transfected with the siRNAs described in Table 5: Per 6-well, 4.4 μL of the respective 10 μM siRNA stock solution was mixed with 5.6 μL of Lullaby transfection reagent (OZ Biosciences, LL70500) in 245.6 μL OptiMEM (Gibco, 31985062) and incubated for 20 minutes at room temperature, before being added dropwise onto the cells. On the following day, the cells were washed, detached, and divided into two separate wells of a 6-well plate. On day 5, the transfection was repeated as described above. The cells were then either harvested on day 6 or day 7 for RNA and protein analysis or seeded 6 hours after the second transfection for transwell migration assays.

**Table 5:**
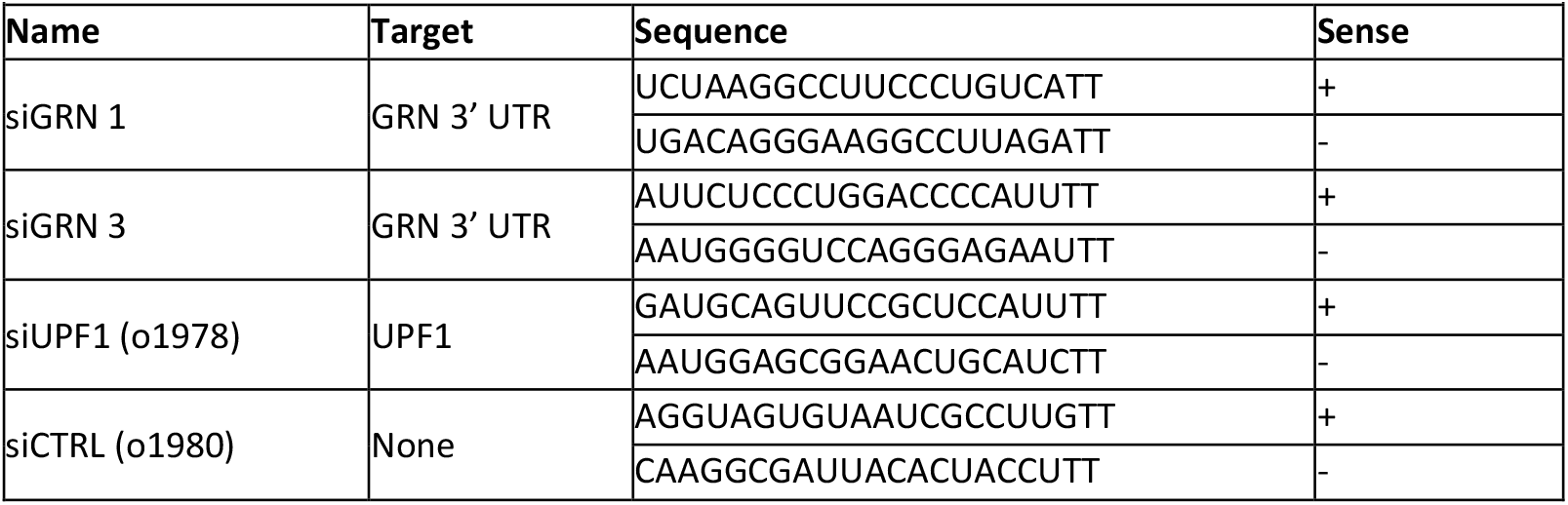
siRNAs.

## Notes

### Competing Interest Statement

The authors have declared no competing interest.

