## Supplemental Figures S1 and S2 for "Analysing the effect of full-length and C-terminally truncated progranulin on proliferation, colony formation, and migration in HepG2 and U87 cells"

**Supplementary Figures S1 and S2** for Hofer et al., *Analysing the effect of full-length and C-terminally truncated progranulin on proliferation, colony formation, and migration in HepG2 and U87 cells*

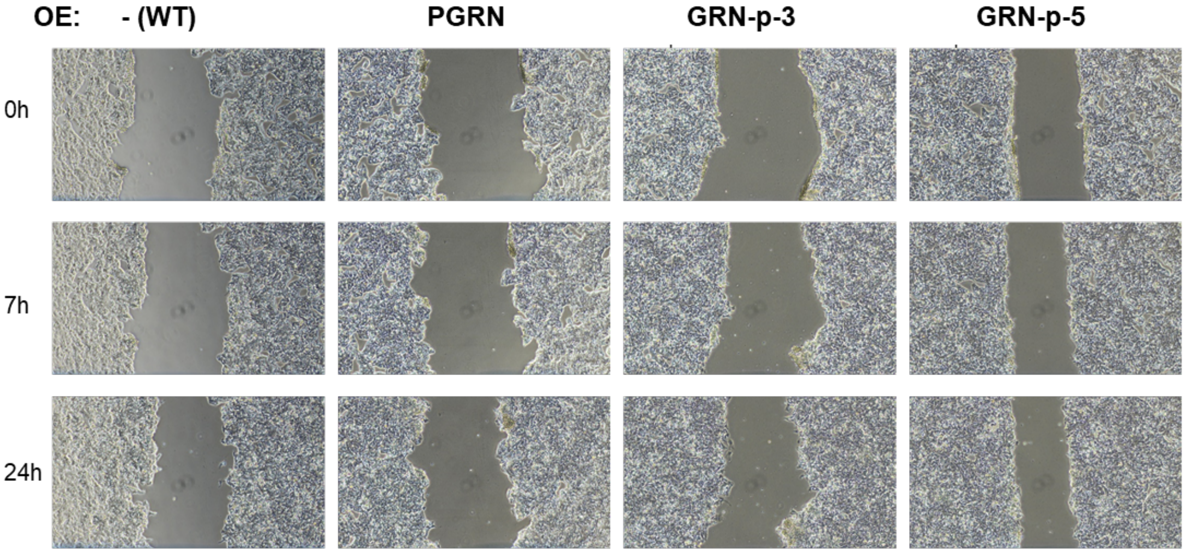


***Supplementary Figure S1: Scratch (wound healing) assay of HepG2 cells**. Cells were grown to high confluency, and then a gap was scratched softly with a pipette tip, which was monitored for 24 hours.*

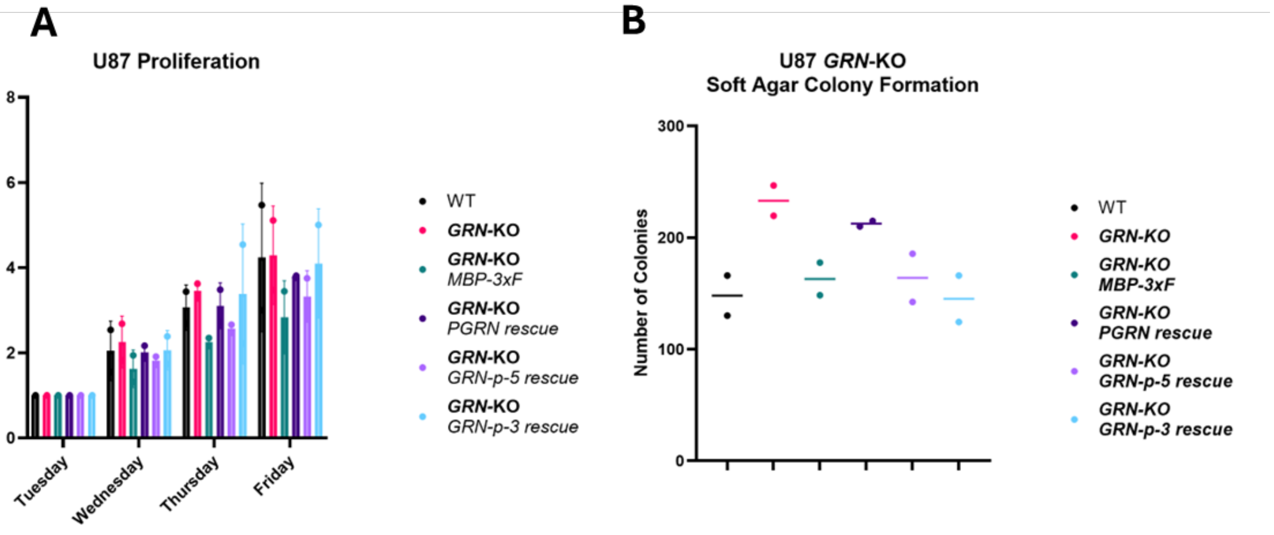


***Supplementary Figure S2: GRN-KO and PGRN-rescue in U87 cells.*** *(A) Proliferation of U87 GRN-KO and rescue cell lines and (B) Soft-agar colony-formation assay (n=2). Panels were created with GraphPad Prism.*
